# HERV-derived epitopes represent new targets for T-cell based immunotherapies in ovarian cancer

**DOI:** 10.1101/2024.07.13.603392

**Authors:** Paola Bonaventura, Olivier Tabone, Yann Estornes, Audrey Page, Virginie Mutez, Marie Delles, Sarah Moran, Clarisse Dubois, Marjorie Lacourrege, Dina Tawfik, Ema Etchegaray, Adrian Valente, Rasha E. Boulos, Gabriel Jimenez Dominguez, Nicolas Chuvin, Nicolas Gadot, Qing Wang, Jenny Valladeau-Guilemond, Stéphane Depil

## Abstract

**Background:** Ovarian cancer represents the most lethal gynecological cancer with poor results of checkpoint inhibitors. Human endogenous retroviruses (HERVs) are aberrantly expressed by tumor cells and may represent a source of shared T cell epitopes for cancer immunotherapy regardless of the tumor mutational burden.

**Methods:** A transcriptomic analysis based on RNA-sequencing (RNA-seq) was developed to quantify the expression of HERV-K sequences containing the selected epitopes. The presence of HERV-K/HML-2 Gag antigen was then assessed by immunohistochemistry (IHC) on tumor microarrays from ovarian cancer samples and normal ovarian tissues. A specific immunopeptidomics approach was developed to detect epitopes on HLA molecules. Epitope-specific CD8^+^ T cells were quantified by multimer staining and *in vitro* target cell killing was evaluated using xCELLigence technology. *In vivo* antitumor efficacy of HERV-specific T cells was assessed in an avian embryo model.

**Results:** Epitope-containing HERV transcripts were significantly higher in ovarian cancers compared to normal tissues. The presence of HERV-K/HML-2 Gag antigen was confirmed by IHC in 20/40 (50%) ovarian cancers while no Gag expression was found in normal ovarian tissue samples. Immunopeptidomics analysis showed the presence of epitopes on HLA molecules on the surface of ovarian tumor cell lines but not on normal primary cells from critical tissues. HERV-specific T cells were detected among tumor infiltrating lymphocytes (TILs) from ovarian cancers, confirming the immunogenicity of these epitopes in patients. *In vitro*, HERV-specific T cells specifically killed ovarian cancer cells in an HLA class I-restricted manner while sparing normal HLA-A2-positive primary cells derived from critical tissues. Epitope-specific CD8^+^ T cells exhibited a strong anti-tumoral activity *in vivo*, inducing a highly significant decrease in tumor volume in comparison with control groups.

**Conclusion:** These results provide the preclinical rationale for developing T-cell based approaches against HERV-K-derived epitopes in ovarian cancer.

**Graphical abstract:** 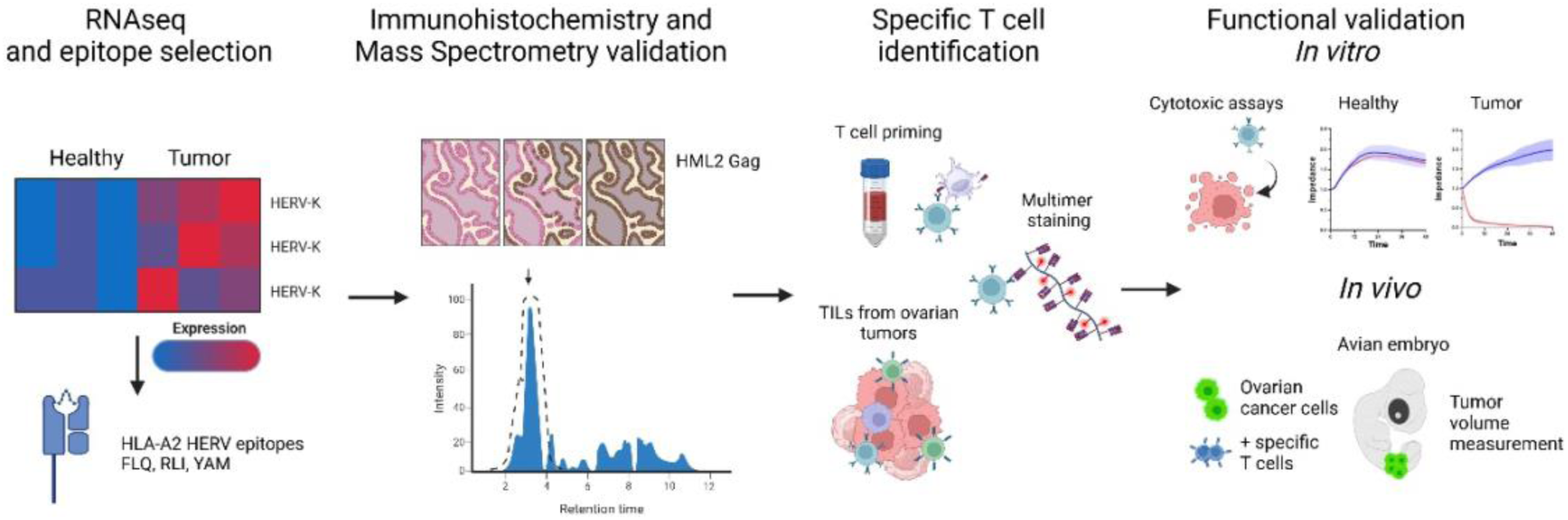

- Some HERVs are specifically overexpressed in ovarian cancer compared to normal tissues.
- HERV-K/HML-2 Gag antigen is detected by immunohistochemistry in ovarian cancers but not in normal ovarian tissues. Furthermore, HERV-K-derived epitopes are presented on HLA molecules on the surface of ovarian cancer cells but not on normal cells.
- These epitopes are immunogenic in patients and induce high-avidity CD8^+^ T cells that specifically kill ovarian cancer cells *in vitro* and *in vivo* while sparing normal cells.

## Introduction

Ovarian cancer represents the most lethal gynecological cancer. The majority of patients are diagnosed at an advanced stage and relapse after first-line chemotherapy is common, leading to poor outcomes [1,2]. Reported results of PD1 or PD-L1 blocking antibodies have been disappointing with less than 10% of objective responses [3]. Furthermore, combinations of anti-PD-L1 antibodies with chemotherapy have also shown negative results [4]. New approaches are thus warranted to induce efficient anti-tumor immune responses in patients with ovarian cancer.

The efficacy of tumor-specific T cells relies on the recognition of tumor epitopes presented on Human Leucocyte Antigens (HLA) molecules on the surface of cancer cells [5]. In this context, neoepitopes derived from non-synonymous somatic mutations have been shown to induce high-avidity tumor-specific T cells [6]. However, the development of personalized cancer vaccines or adoptive T cell therapies targeting this type of epitopes remains challenging, especially in tumors characterized by a low mutational burden, such as ovarian cancer [7]. Thus, there is still an unmet need to identify new tumor epitopes, ideally shared across patients, in ovarian cancer.

Human endogenous retroviruses (HERVs) represent 8% of the human genome. These DNA sequences are the legacy of multiple ancient germ line infections by exogenous retroviruses [8–10]. HERVs are mainly localized in heterochromatin and repressed by epigenetic silencing in normal cells, even if a transcriptional background has been described in normal tissues [11]. Aberrant expression of HERVs has been reported in different tumors, including breast [12], ovarian [13], prostate [14] and kidney cancers [15], melanoma[16], glioblastoma [17], as well as hematological malignancies [18]. Some HERVs, especially the most recently integrated ones (such as HERV-K family), keep partially conserved open reading frame (ORF) encoding Gag, Pol or Env proteins. We have previously demonstrated that HERVs represent a source of shared tumor epitopes capable of inducing T cell clones with high functional avidity. We have further characterized CD8^+^ T cell clones specific to HLA-A2-restricted 9-mer epitopes derived from HERV-K and exemplified their anti-tumor activity in triple negative breast cancer and acute myeloid leukemia [19,20]. Combining a new transcriptomic quantification with immunohistochemistry (IHC) and targeted immunopeptidomics, we report here that HERV-K-derived epitopes are specifically expressed in ovarian cancer cells compared to normal cells. We also demonstrate that HERV-specific T cells selectively kill ovarian cancer cells, both *in vitro* and *in vivo,* while sparing normal cells, providing the preclinical rationale for developing T cell-based approaches against these new targets in ovarian cancer.

## Material and methods

### In silico analyses

#### Datasets

RNA sequencing reads from TCGA were obtained from the GDC portal. They were downloaded in the bam format (already aligned) and were turned into fastq (raw reads) by using the PicardTools software’s SamToFastq (v2.26.10) command. RNA sequencing reads from GTEx V8 (https://gtexportal.org/home/) were obtained using the Terra portal as indicated on their website and were turned into fastq in the same way as TCGA.

#### Tissue selection

From TCGA, we selected all available ovarian tumor samples (n=421) and selected peritumoral healthy samples from the 7 critical tissues with at least 10 samples: Lung (n=110), Kidney (n=129), Liver (n=49), Bladder (n=19), Stomach (n=36), Head and Neck (n=44) and Colon/Rectum (n=51). From GTEx, we selected all the available samples of the same tissue types obtained from TCGA allowing batch correction and we also added other critical normal tissues that were not available in TCGA.

#### Reference annotation

We used the UCSF hg38 as reference genome and Gencode v33 for the gene annotations. For HERV annotation, we extracted the retro.hg38.v1 version from Telescope database [21]. The two annotations were then merged. To avoid duplicates, we ran gffcompare [22] to identify overlaps between regions. We then computed the Jaccard similarity between regions that overlapped; regions with a similarity above 0.5 were considered duplicates, and to keep a consistent nomenclature of HERVs, only the ones originating from the Telescope annotation were kept. We then only considered protein coding genes and HERV transcripts.

#### Transcript quantification

Similarly to SalmonTE [23]and REdiscoverTE [24] pipelines, Salmon [25] (v1.10.0) was used on the unaligned reads to quantify how many reads were associated to each locus. [22] The reference index was built with the index option set to 15, and the reference sequences were generated by using the gffread (v0.12.8) (22) package. Mapping and quantification were then performed by calling salmon count with –validateMappings –l A options.

#### Expression normalization

Obtained pseudo counts for all samples were loaded into R (4.1.2) using tximport function, from tximport library [26]. Before any normalization, we computed for each sample the sum of counts of all HERV loci with a potential Open Reading Frame (ORF) containing the epitope of interest (FLQ, RLI and YAM). These epitope-containing HERV transcript counts were added to the whole dataset, as with any transcript. The counts were then normalized using DESeq2 median of ratio and transformed into log scale with vst function [27].

#### Data exploration and analyses

All analyses were done using R (4.1.2) and RStudio server (2023.12.1-402). Differential Expression analyses between ovarian tumor and all selected peritumoral tissues were performed with DESeq2 (Wald test). P-values were corrected for multiple testing using the Benjamini–Hochberg method. Batch corrections between TCGA and GTEx were run using Combat function [28] from sva package, with TCGA as reference Batch and were done tissue per tissue as previously described [29] Visualizations were done with ggplot2 [30] and ggbiplot library.

#### Peptide sequences matching with Uniprot

The whole reviewed Human Uniprot database (Swiss-Prot) was downloaded from uniprot.org (release 2024_01). Exact sequence pattern matching was perform between each peptide sequence and Uniprot db, and no match was found with any known human protein except with HERV-K-derived proteins.

### Immunohistochemistry

The OV1004a ovary tissue microarray (TMA) from TissueArray was selected for IHC analysis. This TMA contains a total of 40 ovarian tumor samples, each of them in duplicate (supplementary Table 2). Five normal ovarian tissues were also analyzed (from OV1004a FDA3331a, TissueArray;#T8234701-5, BioChain).

IHC was performed on an automated immunostainer (Ventana Benchmark Ultra, Roche, FR) using UltraView Universal DAB detection Kit. Slides were deparaffinized at 72°C using Ventana EZ Prep reagent (Roche, Cat. 950-102) and hydrated, followed by an antigen retrieval method using Ventana Tris-EDTA buffer pH 7.8 (Roche, Cat. 950-224) for 32 minutes at 95°C. Sections were incubated with the HERV-K/HML-2 Gag monoclonal antibody (diluted at 1:200, Austral Biologicals, Cat. HERM-1841-5) for 32 minutes and then with the ultraView Universal HRP Multimer. As a negative control, the microarray was processed similarly but incubated with the ultraView Universal HRP Multimer only. Staining was revealed with 3,3′-diaminobenzidine as a chromogenic substrate for 8 minutes. Then, the sections were counterstained with Gill’s Hematoxylin (Roche, Cat. 760-2021) for 8 minutes and post counterstained with Bluing reagent (Roche, Cat. 760-2037) for 4 minutes.

For staining quantification, TMA were scanned with panoramic scan II (3D Histech, HU) at 20×. Images were generated using software Caseviewer (3Dhistotech 2.4.0.119028) for coloration and IHC. Captured images were processed using HALO AI software (Indica Labs, V3.6.4134) and adapted algorithms for quantification (Multiplex IHC v3.4.9). Each spot was annotated by an anatomopathologist to outline the tumor and eliminate artefact zones. The Mininet neural network was used to train the software to discriminate stroma and tumor structures. The nuclear and cytoplasmic stains were quantified with an algorithm taking into account the percentage of positive cells per spot and cell staining intensity (three intensities used: low, moderate, high). Staining was also evaluated by two operators and scored manually (scores: 0, 1, 2 or 3) according to staining intensities. The final quantification considers both HALO AI software quantification and scores obtained by both operators. A score of 3 was classified as high, scores of 2 or 1 as moderate and score of 0 as negative.

### Epitope detection by mass spectrometry (MS)

Epitope validation by MS was performed by Complete Omics Inc. (Maryland, USA) according to the method previously described [31] with additional modifications. In brief, for sample preparation, at least 20 million cells were lysed and peptide-HLA complexes were isolated using in-house packed Valid-NEO neoantigen enrichment column preloaded with anti-human pan-HLA antibodies modified with conjugated chemical moieties that are designed to increase the binding efficiency between the column matrix and HLA-epitope complex molecules. MaxRec technology is conducted to increase the sensitivity of the detection by preventing sample loss. Several versions of MaxRec peptides were developed to compare the difference in protection efficiency, together with chemically modified MaxRec peptides. After elution, dissociation, filtration and cleanup, epitope peptides were lyophilized before further analysis. Detection parameters for each epitope were examined and curated through Valid-NEO method builder bioinformatic pipeline developed by Complete Omics Inc. Precursor or fragmented ions with excessive noise due to environment or coelution with impurities were excluded from the detection. To boost up the detectability, a series of recursive optimizations for the significant ions were conducted.

### Biological samples

Blood from healthy donors was obtained from the ‘‘Etablissement Français du Sang’’ (Lyon, FR). Fresh high-grade serous ovarian carcinomas (n=13) were provided by the tissue bank of Centre Léon Bérard (CLB) (BB-0033-00050, CRB - CLB, Lyon, France; French agreement number: AC-2013-1871), after approval from the institutional review board and ethics committee (L-06-36 and L-11-26) and patient written informed consent, in accordance with the Declaration of Helsinki.

### Cell lines and primary cell cultures

T2 (Cat. ACC 598) cells were purchased from DSMZ (GE), OVCAR-3 (Cat. HTB-161) cells were purchased from ATCC (US) and cultured according to the manufacturer’s instructions. 1% penicillin-streptomycin (PS, Gibco Cat. 15140122) was added to the culture medium. HLA-A2^+^ normal primary cells HCM (Cat. C-12810, batch#463Z016.2, female donor DN000249), HBEpC (Cat. C-12640, batch#469Z016, female donor DN000331) and NHEK (Cat. C-12003, batch#467Z005.1, female donor DN000068) were purchased from Promocell GmbH (GE), nHKPT cells (Cat. 3253021, batch#RPCT022823, male donor) from Tebubio (FR) and HA cells (Cat. 1800-SC, batch#35382) from ScienCell (US). All human normal primary cells were cultured following the supplier’s recommendations. All cells were grown at 37°C in 5% CO2 with humidification. OVCAR-5 tumor cell lines (NCI-60 panel) for MS-immunopeptidomics analysis were provided by Complete Omics Inc.

### Fresh tumor dilaceration and expansion of tumor infiltrating lymphocytes (TILs)

Tumor tissues were dissected into fragments of approximately 1 mm^3^ and digested with collagenase IV (Sigma, Cat. C2674) and DNAseI (Sigma, Cat. D4513) during mechanical dilaceration, for 45 minutes in 20% SVF (Eurobio, FR Cat. CVFSVF00-01, batch S80412-4602) supplemented RPMI. The tumor lysate was centrifuged at 1500 rpm for 5 minutes, counted and resuspended in 5% human serum enriched RPMI (Gibco Cat. 52400041). Cells were counted and plated at a density of 5×10^4^ cells per well in a flat bottom 96-well plate with anti-CD3 anti-CD28 Dynabeads (Dynabeads™ Human T-Expander CD3/CD28 Cat. 11141D, Gibco,FR) and IL-2 (Proleukin, Vidal Cat. at 100 IU/mL) in a ratio beads to cells of 1:4, to expand TILs. Dextramer staining was performed on TILs expanded for 14 days after tumor dilaceration. Two to five hundred thousand cells per condition (epitope specificity, or controls) were washed in 2 mL washing buffer (PBS + 2% FBS + 2mM EDTA (Sigma Alderich, Cat. E7889 US)) and stained for 10 minutes with dextramers (Immudex ApS, DK) at room temperature prior to viability and surface marker staining and then washed two times. A dextramer complexed to a non-natural irrelevant peptide (ALIAPVHAV, HLA-A2 neg. control Cat. WB02666 Immudex ApS, DK, EU) was used as negative control.

All samples were analyzed on a LSR-Fortessa (BD Biosciences, FR) with calibrated settings to compare samples throughout the entire study. Data were analyzed using FlowJo Software (Tree Star v10.4, NJ, USA).

### *In vitro* priming assays and generation of epitope-specific CD8^+^ T cells

Peripheral blood mononuclear cells (PBMCs) were obtained by Ficoll density gradient centrifugation (Eurobio, FR,Cat. CMSMSL01-01). Monocyte were isolated and cultured for 6 days in RPMI medium in the presence of GM-CSF (Peprotech, Cat.300-03) 200ng/ml and IL4 (Peprotech, Cat. 200-04) 50ng/ml for the differentiation of monocyte-derived dendritic cells (MoDCs). MoDCs were matured with TNF-a (Peprotech Cat. 300-01A 20ng/ml) and PolyIC (Invivogen Cat. tlrl-PIC-5 40µg/ml) 16 hours before pulsing with the peptide of interest and cultured with autologous T cells in a MoDC:T ratio of 1:10. Cells were co-cultured for 14 days in AimV medium enriched with IL-7 (Miltenyi Cat.130-095-363) and IL-15 (Peprotech Cat.200-1s), changing media periodically and restimulating a second time with peptide-pulsed MoDCs after 7 days. At the end of the co-culture cells were harvested and counted for tetramer staining analysis. Tetramers were produced by conjugate biotinylated HLA-A*02:01 peptide-specific monomers (P2R Facility, Nantes) with PE-Streptavidin (Biolegend, US, Cat. 405204) or APC-Streptavidin (Biolegend, Cat.405207). Specific cells were isolated by magnetic monomer sorting. Positive and negative fraction were cultured separately and expanded on feeders composed by 35 Gy-irradiated allogeneic PBMCs and B-lymphoblastoid cell lines (kindly provided by Dr. Henri Vie CRCI2NA-Equipe12 INSERM UMR 1307 / CNRS UMR6075). Feeder cells were plated in a 96-well round bottom plate at a concentration of 0.10×10^6^ cells per well in RPMI 5% human serum with PHA-L 1.5µg/mL (Merck KgAa, GE, Cat.30852801), IL-2 150 IU/mL and IL-7 (10ng/ml). Cells were cultured for 14 days, and medium was replaced when needed with IL-2 and IL-7 enriched fresh RPMI 5% human serum. This process was repeated if needed to obtain a specific population (>80%).

### T cell functional avidity and IFNγ ELISPOT

Functional avidity of specific CD8^+^ T-cells was assessed using IFN-γ ELISPOT assay (IFNγ ELISPOT, Diaclone, FR, Cat.856-051-020). Epitope-specific CD8^+^ T cells were restimulated with T2 cells pulsed with decreasing concentrations of the cognate peptide (from 10^-6^ to 10^-13^ M) or with an irrelevant peptide (10^-6^ M) used as negative control in a T cell: T2 cell ratio of 1:10 in RPMI medium (Gibco, FR, EU) supplemented with 8% of human serum. After 18 hours, supernatants were removed and ELISPOT was performed. The minimal peptide concentration required to achieve a significant cytokine response was determined (>30 spots and 2-fold change in comparison to the negative control). The peptide concentration required to achieve a half maximal cytokine response (EC_50_) was determined (graphpad prism, version 6.0 for Windows was used for the 50% EC (EC_50_) determinations, R>0.98).,

### xCELLigence killing assay

Killing assays were performed using an xCELLigence RTCA eSight Cell Analyzer (Agilent, FR). OVCAR3 tumor cell line and normal primary cells were seeded in E-plates 16 (Agilent) and cultured overnight before adding T cells in Effector (E):Target (T) ratio of 2:1 or 5:1. When required, synthetic peptides at a final concentration of 1 nM for 2 hours at 37°C, blocking anti– MHC-I antibody (50 μg per well; clone W6/32, Bio X Cell, Cat. BE0079) or isotype control (final volume 100μl) were added. Annexin V FITC (Biolegend, FR, Cat. 640945) was added to each well (f.d. 1/1000) for cell death analysis in live imaging

Impedance variation (cell index) measurement and live imaging acquisition were performed in real time, every 15 minutes and for 48 hours, after the addition of T cells and each condition was performed in triplicates. The normalized cell index (NCI) was calculated at the time of T cell addition. Percent of lysis was calculated using RTCA Software Pro Immunotherapy Module and normalized on unspecific T cell. Annexin V staining was measured on images (green object count/image) at different time points (0h, 6h, 24h and 48h after the beginning of the co-culture) using the RTCA eSight software.

## In *vivo* anti-tumor efficacy

*In vivo* anti-tumoral activity of HERV-specific CD8^+^ T cells was performed at Oncofactory (Lyon, FR). Embryonated eggs were obtained from a local supplier (EARL Les Bruyeres, FR). Laying hens’ sanitary status was regularly checked by the supplier according to French laws. Development of embryos and *in ovo* xenografts were performed as previously described [32]. Eggs were incubated at 38.5°C in a humidified incubator until the desired developmental stage (stage HH14, 2 days post-fertilization). Embryos were randomized in each experimental group and were harvested at embryonic day 4 (stage HH25, 4 days post-fertilization).

OVCAR-3 cells were labelled with 7 µM CFSE solution (Life Technologies, Cat. C34554) and mixed with T cells at a E:T of 5:1 before co-engraftment. At stage HH14 chick embryos were co-engrafted with mixed cells in presumptive somitic areas with a glass capillary connected to a pneumatic PicoPump (PV820, World Precision Instruments) under a fluorescence stereomicroscope. Targeted tissue areas for the graft were visualized under the stereomicroscope. Co-transplanted embryos were harvested after 48 hours (at stage HH25) and *in vivo* anti-tumoral effect of the CD8^+^ T cells was evaluated by assessing the tumor volume using 3D light-sheet imaging. For this analysis, tissues from n=45 chicken embryos were cleared and imaged using whole-mount SPIM imaging with volumetric analyses performed as previously described [32].

Data are presented as mean ± s.d. and one-way ANOVA post-hoc Dunnett’s test was used for multigroup comparisons (Fig.6B and Supplementary Fig.5B). P values were considered statistically significant if P<0.05, with ****P<0.001.

## Results

### HERV-K antigens are selectively expressed in tumor cells compared to normal cells

Based on our previous study [19], we selected 3 HLA-A2-restricted epitopes derived from cancer-associated HERV-K sequences, which were shown to be immunogenic and able to induce high-avidity cytotoxic T cells. The two 9-mer epitopes FLQFKTWWI (FLQ) and RLIPYDWEI (RLI) derive from HERV-K Gag and YAMSNLFSI (YAM) from HERV-K Pol. To quantify the expression of these epitopes at the transcriptomic level in ovarian cancer, we first identified all the HERVs with a predicted open reading frame (ORF) containing the corresponding epitope sequences. The sequences of FLQ and RLI epitopes were found in 33 and 37 HERVs, respectively, while the sequence of YAM was detected in 43 HERVs (supplementary Table 1). Importantly, these peptides were not found in any known protein sequence in Uniprot except from HERV-K-derived proteins. Differential expression analysis performed between ovarian cancer and critical peritumoral tissues from The Cancer Genome Atlas (TCGA) normalized RNA-seq data (Fig.1A), showed that the majority of the HERV loci containing one or several epitope sequences are more expressed in ovarian tumor samples than in all peritumoral tissues (Fig.1B). In more details, HERV-K/HML-2_19q11 is the most differentially expressed (DE) transcript containing FLQ peptide, HERV-K/HML-2_19q11 and HERV-K/HML-2_11q12.3b are the most DE transcripts containing RLI and HERV-K/HML-3_10q22.1 is the most DE transcript for YAM peptide (Fig.1C). Then, we computed the sum of the raw counts for all the transcripts containing a given epitope sequence in each sample to make comparisons of its expression (Fig.1A). The sum of the epitope-containing HERV transcripts was significantly higher (P<0.01) in ovarian cancer for all 3 peptides compared to the 7 normal tissues considered as critical, with more than 1 log_2_-fold change in most tissues (Fig.1D). To verify that the peritumoral tissues available in TCGA are representative of the normal critical tissues, we also processed normal samples from the Genotype-Tissue Expression (GTEx) RNA-seq database in the same way as for TCGA tissues. The expression of the epitope-containing HERVs in GTEx healthy critical tissues was not higher in brain, blood vessels, blood, heart, skin or muscle than in the other critical tissues also available in TCGA (supplementary Fig.1A). To validate that the epitope-containing HERVs are more expressed in ovary tumors than in normal tissues, we aimed to include GTEx-related samples in addition to the TCGA dataset. As shown by the UMAP and the HERVs expression levels (supplementary Fig.1B and 1C), GTEx and TCGA datasets cannot be compared directly. We therefore run a batch correction on the whole dataset using ComBat [28] for each tissue independently (except for ovarian peritumor samples that are not available in TCGA) (supplementary Fig.1D). This approach confirmed that the expression levels are significantly higher (P<0.01) in ovary tumors than in any normal peritumoral tissue for all 3 peptides (supplementary Fig.1E).

**Figure 1:**
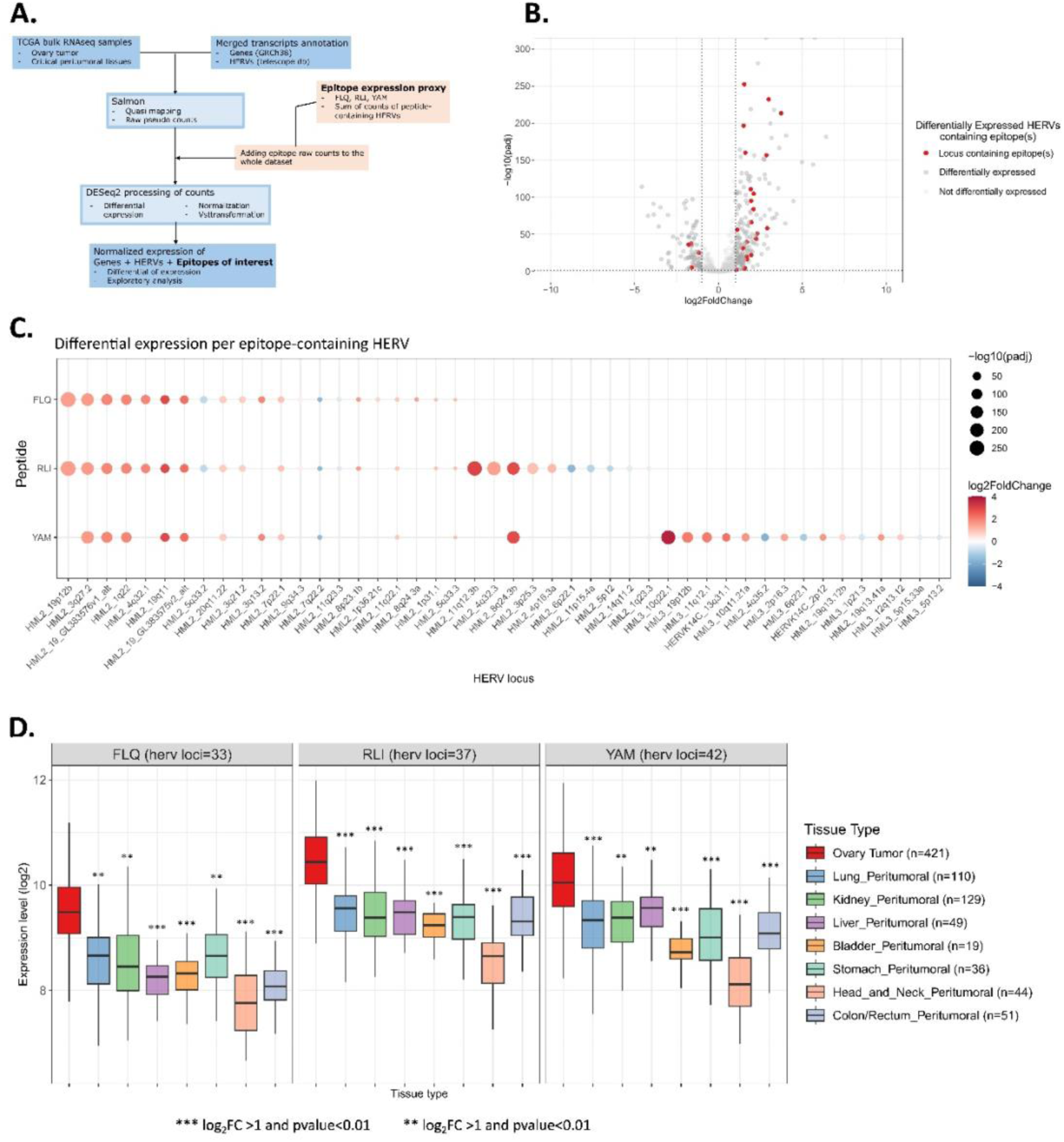
Transcriptomic analysis of HERV epitope expression. **A.** Schematic representation of data processing. Salmon was used to map and quantify reads to a merged reference transcripts annotation containing both genes and HERVs annotations. Transcriptomic expression of the epitope was estimated by the sum of all HERV loci with a potential ORF containing its sequence (FLQ, RLI, and YAM). Data were normalized and log transformed (vst) using DESeq2; ORF: open reading frame. **B.** Volcano plot showing differentially expressed HERV-K loci in ovary tumor against peritumoral tissues from TCGA. Dark-grey dots represent the HERV-K loci differentially expressed (log_2_FC>1 and FDR<0.01, Wald test from DESeq2). Red dots represent the HERV-K loci that are differentially expressed and contain FLQ, RLI and/or YAM epitopes in a potential ORF; FC: fold change; FDR: false discovery rate. **C.** Differential expression of epitope-containing HERV loci in ovary tumor against peritumoral tissues from TCGA. Red and blue indicate respectively over- or under-expressed HERV loci in ovary tumor compared to peritumoral tissues; color intensity represents the degree of perturbation (log_2_FC) and size of dots the statistical significance (FDR). **D.** Epitope-containing HERVs in ovary tumor (red) and peritumoral tissues from TCGA. Epitope expression levels represent the sum of epitope-containing HERVs normalized and log_2_ transformed (vst) using DESeq2. The number of HERV loci containing each epitope is represented on top (HERV loci). The number of samples for the corresponding tissues are indicated in the legend (n). Expression levels in ovary tumor were compared to each tissue for all 3 peptides. (*** log_2_FC >1 and p-value<0.01, ** log_2_FC >0.75 and p-value<0.01, * log_2_FC >0.5 and p-value<0.01, Wilcoxon test).

To determine whether this transcriptomic expression is associated with protein translation, we performed IHC experiments using an HERV-K/HML-2 Gag antibody on TMA of ovarian cancer samples. Staining intensities were ranked as high, moderate or negative. A Gag staining was observed in 20 out of the 40 (50%) tested samples. Six (15 %) ovarian tumors were ranked as high, 14 (35%) as moderate and 20 (50%) were negative (Fig. 2A). No Gag staining was found in normal ovarian tissue samples (n=5). These results suggest that HERV-K RNA overexpression observed in ovarian cancer is sufficient to induce the translation of a Gag protein detected by classical IHC in some cancer samples.

**Figure 2:**
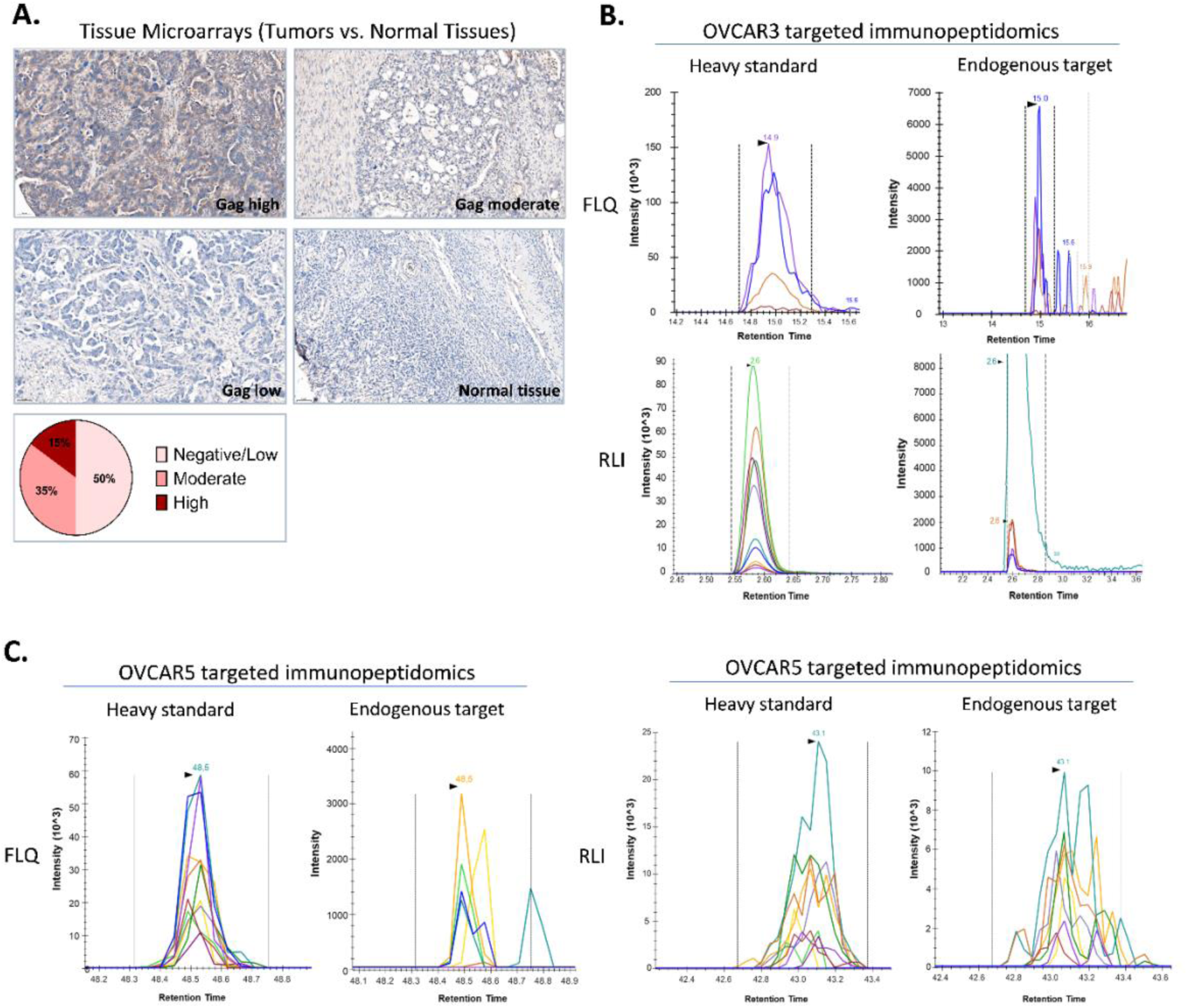
HERV-K antigens are selectively expressed in ovarian cancer. **A.** Immunohistochemistry of HERV-K/HML-2 Gag antigen in tissue microarrays of ovarian cancer and healthy ovarian samples (20x at microscope and 22x for images, images processed with Halo AI software (Indica Labs, V3.6.4134)). Representative examples of tumor and normal ovarian tissues and pie chart of Gag expression level (low/negative, moderate and high) in the 40 tested samples. **B, C.** Targeted immunopeptidomics analysis (Max Rec_Ver2, Complete Omics Inc.) of FLQ and RLI presented on HLA molecules on the surface of OVCAR3 (B) and OVCAR5 (C) ovarian cancer cell lines. Heavy peptides are used as positive control (standard).

We then assessed whether the selected epitopes are efficiently presented on HLA molecules on the surface of tumor cells using a highly sensitive targeted MS-based immunopeptidomics analysis of HLA class I-eluded peptides from two HLA-A2-positive ovarian cancer cell lines, OVCAR-3 and OVCAR-5. The Valid-NEO method which uses heavy isotope-labeled peptides to accurately characterize by MS the presence of endogenous peptides [33], was applied and optimized for FLQ and RLI and allowed the detection of both peptides with high confidence signal (Fig.2B and 2C). A targeted Valid-NEO method for YAM analysis could not be optimized from a first series of experiments, therefore the presence of YAM-HLA complexes on OVCAR-3 and OVCAR-5 could not be confirmed. To validate the tumor selectivity of these epitopes, peptide-HLA complexes were also investigated in 5 HLA-A2-positive primary normal cells from a selection of critical tissues: kidney proximal tubules (NhKPT), cardiomyocytes (HCM), astrocytes (HA), keratinocytes (NHEK) and bronchial epithelial (HBEpC). By using the same method and MS sensitivity, no epitope-HLA complexes were detected on any of the tested primary normal cells (supplementary Fig.2).

### T cells specific for HERV-K epitopes can be detected in ovarian tumors

We next sought to determine whether these HERV-derived epitopes are immunogenic in patients by looking for the presence of specific T cells among TILs from ovarian tumors (Fig.3). Due to the limited numbers of cells obtained from tumor biopsies, we first polyclonally expanded TILs *in vitro* without any addition of peptide before detection of HERV-specific T cells by dextramer staining in flow cytometry (Fig.3A). Dextramer-specific T cells for at least one HERV epitope were detected in 10 of the 13 (77%) analysed samples (including 12 high-grade and one low-grade serous carcinomas). FLQ, RLI and YAM-specific T cells were found in 9/13 (69%), 3/13 (23%) and 7/13 (54%) patients, respectively. A higher staining (>0.05% dextramer-positive CD8^+^ T cells) was observed in 2 patients for the FLQ epitope and 2 other patients for the YAM epitope (Fig. 3B and 3C). These results suggest that the 3 epitopes can induce specific CD8^+^ T cells in patients with ovarian cancer.

**Figure 3:**
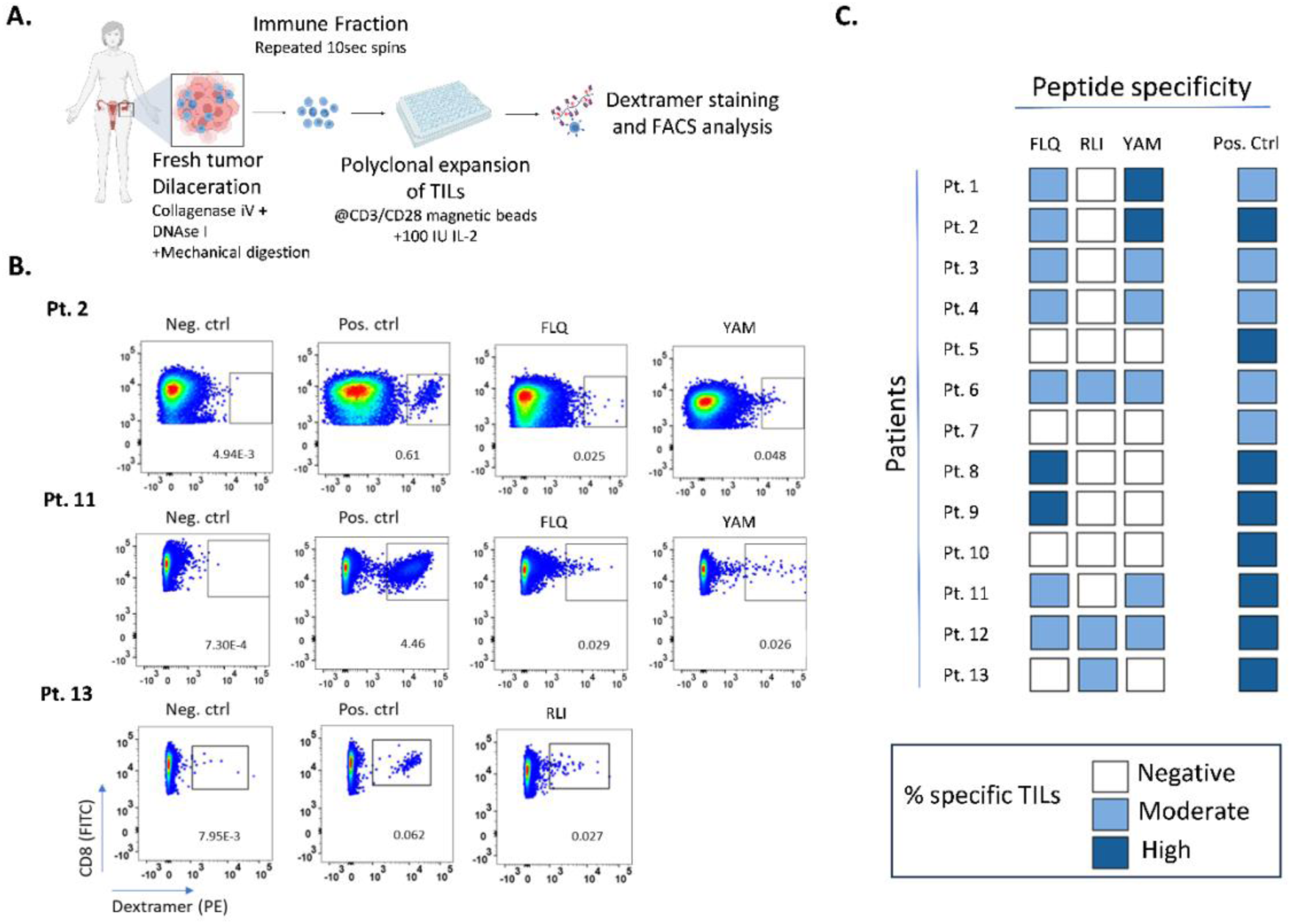
HERV-K-specific CD8^+^ T cells are detected in ovarian tumors. **A.** Schematic representation of tumor and TIL processing before specific dextramer staining **B.** Representative plots of CD8^+^ T cells (Y axis, FITC) with quantification of the percentage of dextramer^+^ cells (x axis, PE, with positive population framed) in polyclonally expanded TILs from ovarian cancer samples. From left to right: dextramer negative control, dextramer positive control and HERV-K epitopes. Pt: patient number. **C.** Summary of the results obtained with dextramer specific staining of TILs derived from ovarian cancer patients’ samples (n=13). White squares represent the absence of a specific staining (negative <0.01% specific TILs), light blue squares represent moderate specific staining (0.01% ≤ x ≤ 0.05% or ≥0.05% with MFI <10^3^), blue squares represent high specific staining (>0.05% specific TILs); pos. Ctrl: positive control.

### HERV-specific CD8^+^ T cells kill ovarian cancer cells while sparing normal cells

Next, we assessed the anti-tumor activity of HERV-K-specific T cells generated from healthy HLA-A2-positive donors. Epitope-specific CD8^+^ T cells were stimulated *in vitro* using autologous dendritic cells pulsed with the cognate peptide and further amplified after immunomagnetic sorting. This approach generates a population of specific T cells containing a predominant clone, as previously described [19] (supplementary Fig.3A-C). We first confirmed that the HERV-specific T cells are characterized by a high functional avidity. Elispot assays showed T cell specific IFNγ secretion after restimulation with T2 cells pulsed with the cognate peptide but not with an irrelevant peptide used as control. Remarkably, IFNγ secretion was still detected at peptide concentrations below 10^-10^ M, in line with EC_50_ estimated at 1.6×10^-10^, 1.5×10^-11^ and 1.9×10^-10^ M for FLQ, RLI and YAM-specific T cells, respectively (Fig.4A and supplementary Fig.3D). We then assessed the capacity of these HERV-specific CD8^+^ T cells to recognize and kill ovarian cancer cells. Tumor cell death was monitored in real time using the xCELLigence RTCA eSight technology. T cell clones induced a specific killing of OVCAR-3 cells, as shown by the rapid decrease of the cell index and the corresponding increase of the percentage of cell lysis, whereas the tetramer-negative fraction of T cells did not show any significant effect on impedance. The effect was inhibited by an anti-HLA class I antibody, confirming that the observed cytotoxicity is HLA class-I dependent (Fig.4B-C and supplementary Fig.3E-F). Annexin V staining was added to the medium and the image analysis was performed together with the impedance analysis to further validate the results. Imaging results showed a time-dependent and HLA-restricted increase of annexin V staining when ovarian cancer cells were co-cultured with HERV epitope-specific CD8^+^ T cells but not control T cells (Fig.4D-E).

**Figure 4:**
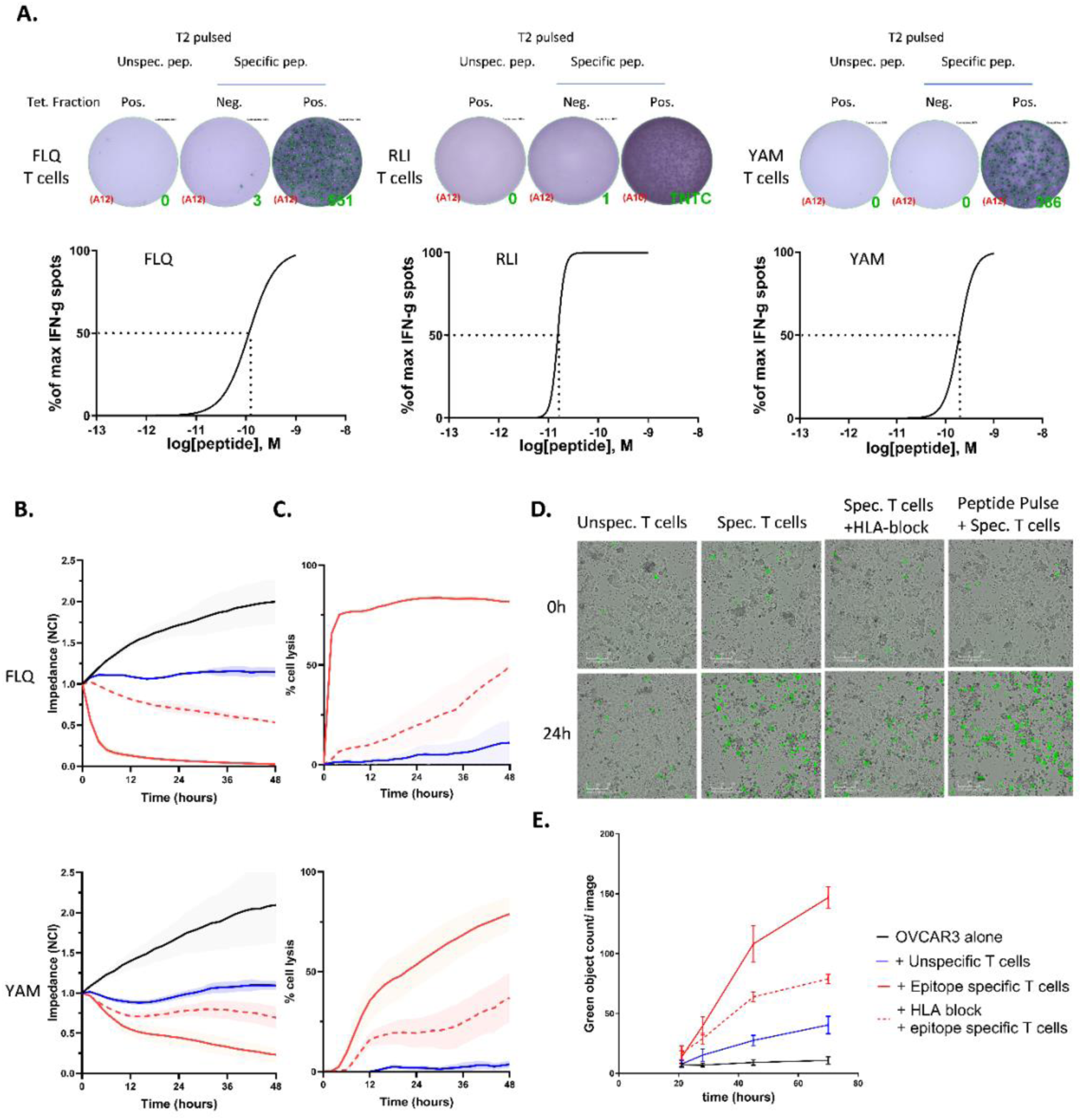
HERV-K derived epitopes induce high-avidity CD8^+^ T cells that kill ovarian tumor cells *in vitro*. **A.** ELISPOT of IFNγ production of tetramer-negative and positive T cell fractions after restimulation with T2 pulsed with irrelevant or cognate peptide (10^-9^M, left panels, one representative well). Nonlinear fit curves of IFNγ spot count per 1 × 10^6^ T cells for specific CD8^+^ T cell clones co-cultured with irrelevant-pulsed T2 or T2 pulsed with a limiting dilution of the cognate peptide. Results are mean of n=4 for FLQ and YAM and n=2 for RLI (independent experiments). EC_50_ is determined using nonlinear fit of mean IFNγ spot count (GraphPad); tet: tetramer; unspec.: unspecific; pep: peptide; pos.: positive; Neg.: negative; TNTC: too numerous to count. **B and C.** Representative plots of tumor cell impedance (cell index) (**B**) or percentage of cell lysis(**C**), of OVCAR3 cells in co-culture with FLQ-(upper quadrants, E:T ratio 2:1), YAM-(lower quadrants, E:T ratio 5:1) specific CD8^+^ T cells, or the corresponding tetramer-negative fractions. Results are mean ± s.d. of technical triplicates of a representative experiment of n=3 for FLQ and n=2 for YAM; E:T: effector to target. **D.** Representative 10X images of co-cultures of FLQ-specific T cells with OVCAR3 presented in C and D at 0h and 24h. Annexin V^+^ dead cells are depicted in green; unspec.: unspecific; spec.: specific. **E.** Cell death quantification of annexin V^+^ OVCAR3 cells (green object/image) at 0, 6, 24 and 48h after the addition of FLQ-specific CD8^+^ T cells or unspecific T cells. Results are expressed as mean ± s.d. of 2 different images In **C, D, F**, target cells alone are depicted with black lines, HERV epitope-specific T cells are depicted with red lines, unspecific T cells are depicted with blue lines. Addition of the anti-HLA antibody to co-cultures is depicted with the red dotted line. NCI : normalized cell index

We next analyzed the potential cytotoxicity of the specific T cell clones toward the panel of HLA-A2 primary normal cells derived from normal critical tissues. In agreement with the absence of epitope presentation on HLA molecules, we did not observe any significant effect of HERV-specific T cells against normal cells, confirming the tumor selectivity of these epitopes (Fig.5 and supplementary Fig.4).

**Figure 5:**
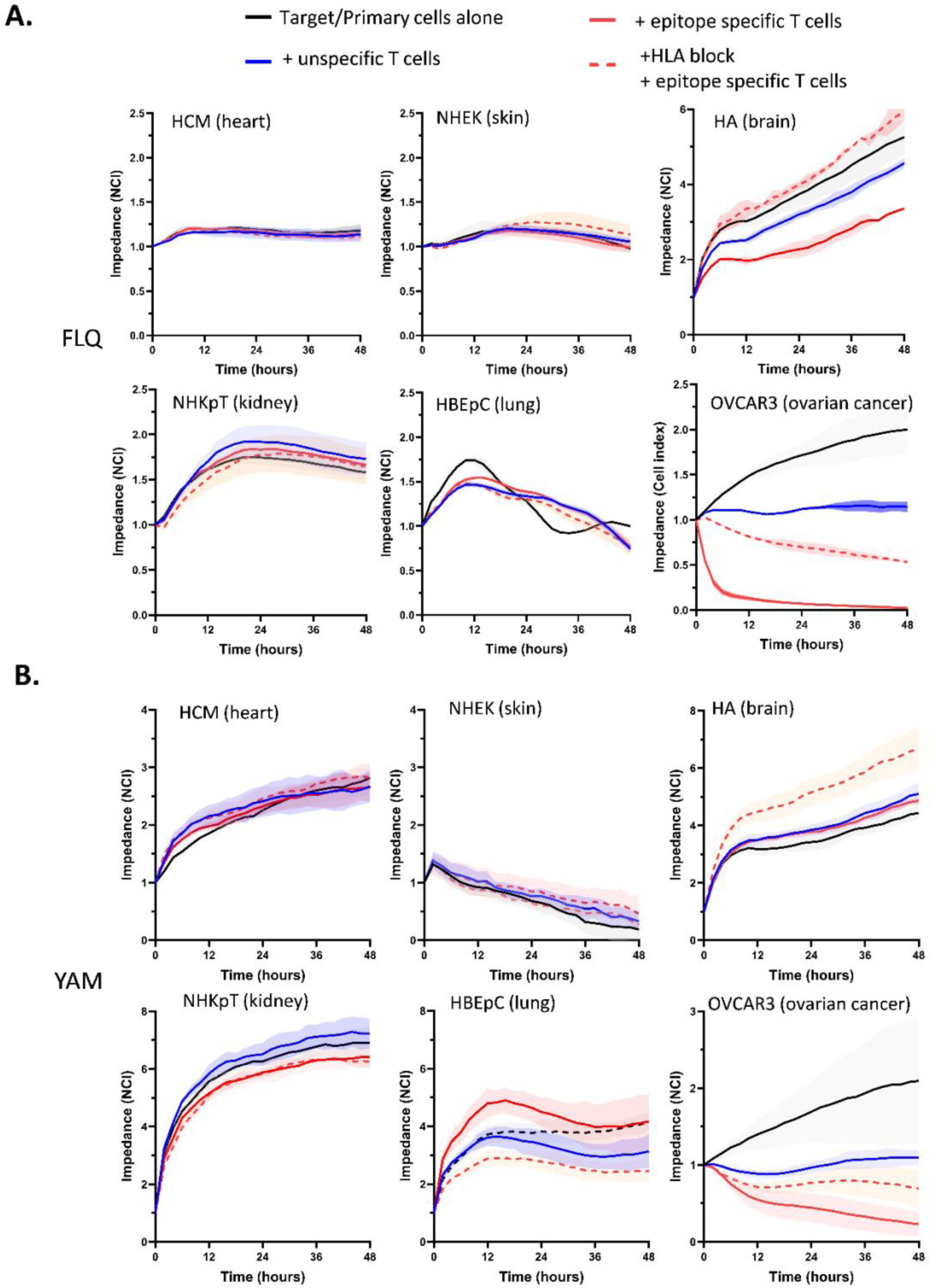
HERV epitope-specific T cells are not cytotoxic toward normal primary cells. **A.** Viability of normal primary cells (HCM, NHEK, NHEK, HA, NHKpT, HBEpC) alone or co-cultured with unspecific T cells (negative tetramer fraction) or FLQ-specific T cells (5:1 effector to target cell ratio) monitored by normalized cell index through xCELLigence. **B.** Viability of normal primary cells (HCM, NHEK, NHEK, HA, NHKpT, HBEpC) alone or co-cultured with unspecific T cells (negative tetramer fraction) or YAM-specific T cells (5:1 effector to target cell ratio) monitored by normalized cell index (NCI) through xCELLigence. OVCAR-3 cells were used as positive control. Results are mean ± s.d. of technical triplicates. HCM, human cardiac myocytes; NHEK, normal human epithelial keratinocytes; NHEK, normal human epithelial keratinocytes; NHKpT, normal human kidney proximal tubule cells; HBEpC, human bronchial epithelial cells. NCI : normalized cell index

### In vivo antitumor efficacy of HERV specific T cells

We then evaluated the *in vivo* anti-tumor efficacy of HERV-specific CD8^+^ T cells. Here, we used the avian embryo tumoral model that consists in implanting human cancer cells within selected embryonic tissues and allows to establish 3D tumors in a few days [32,34] (Fig. 6A). FLQ-specific CD8^+^ T cells co-transplanted with the ovarian cancer cell line OVCAR-3 exhibited a strong anti-tumoral activity, inducing a highly significant decrease in tumor volume in comparison with both control groups in which tumor cells were transplanted alone or with polyclonal T cells (Fig. 6B-C). Indeed, a mean decrease of 63% (P<0.001) in normalized tumor volumes was observed in embryos transplanted with the epitope-specific T cells, as compared to the control group transplanted with polyclonal unspecific T cells (Fig. 6B and supplementary Fig. 5A-B). These latter results confirmed that HERV-specific T cells recognize and kill ovarian cancer cells *in vivo* in a 3D tumor environment.

**Figure 6:**
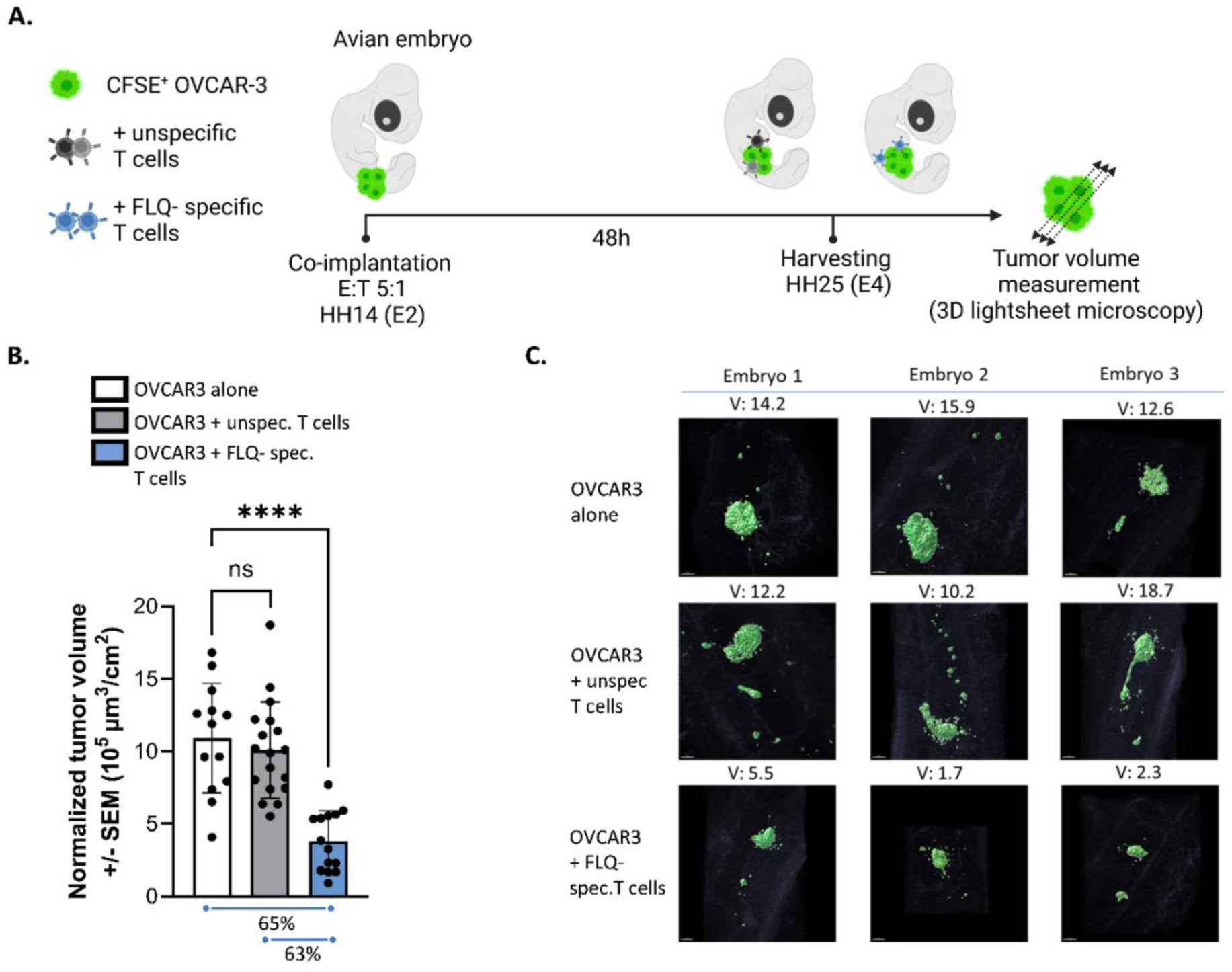
HERV-K epitope-specific T cells elicit *in vivo* anti-tumor effects. **A.** Schematic representation of the *in vivo* experiments. Effector to target (E:T) cell ratio used 5:1; HH, Hamburger-Hamilton stage; E, embryonic day. **B.** Quantification of OVCAR-3 normalized tumor volumes using 3D light sheet microscopy of fluorescent target cells, after 48h co-engraftment with FLQ-specific CD8^+^ T cells (E:T = 5:1). Tumor volumes are normalized against body surface area (BSA). Results are mean ± s.d. of 45 embryos (target cell alone n=13; unspecific T cells n=18; FLQ-specific CD8^+^ T cells n=14); unspec.: unspecific; spec.: specific. ****P<0.001, one-way ANOVA post-hoc Dunnett’s test. **C.** Representative images and volumes for 3 representative embryos at 48h of 3D views (light-sheet imaging) HH25 chick embryos co-engrafted with CFSE-labelled OVCAR3 alone (upper panels) and unspecific T cells (central panels) or FLQ-specific CD8^+^ T cells (lower panels); unspec.: unspecific; spec.: specific; V: tumor volume (10^5^ μm^3^/cm^2^).

## Discussion

HERV expression has previously been reported in ovarian cancer. Pioneering works in the field were published by Wang-Johanning *et al*, showing the presence of HERV-K Env protein in ovarian cancer and the possibility to generate T cells that are cytotoxic against tumor cells using dendritic cells transfected with *Env* [13,35]. Another paper recently published confirmed the expression of HERVs in ovarian cancer and proposed an HERV prognostic score associated with survival and correlated with T cell infiltration [36]. Consistent with these results we demonstrated here that HERV-K-derived epitopes are selectively expressed and presented on HLA molecules on the surface of ovarian tumor cells, but not on normal cells from critical tissues. We also validated the cytotoxicity of high-avidity HERV-specific T cells against ovarian cancer cells, both *in vitro* and *in vivo*.

The transcriptomic quantification of these epitopes is challenging because the sequence of each epitope can be found in independent transcripts from different HERV loci. In this work, we used an approach based on the sum of all the transcripts containing a potential ORF encoding the epitope, as previously described [37]. This method can be considered as one of the most accurate to quantify HERV expression at family level [38]. Doing this, we confirmed that the epitope-containing HERVs are significantly overexpressed in ovarian cancer compared to normal tissues. Importantly, our study highlighted the bias existing when directly comparing datasets from different sources and databases (batch effect). Indeed, HERV counts are systematically higher in GTEx than in TCGA for comparable tissues. As a consequence, we performed a batch correction using ComBat [28] for each tissue available in both TCGA and GTEx databases, as previously reported [29]. We used TCGA as the reference batch, so that overall expression values for each corresponding tissue from GTEx aligned with TCGA samples. After pooling all the corrected batch samples together, we added expression values from ovary tumor samples to compare with normal tissues. These results confirmed that the HERVs containing the RNA sequences coding for the 3 peptides are significantly overexpressed in ovarian cancer compared to normal tissues.

Importantly, we showed here that the overexpression of epitope-containing HERVs at the transcriptomic level is associated with the presence of HERV-K/HML-2 Gag protein in some ovarian cancer samples and with a selective epitope presentation on ovarian cancer cell lines. This supports our hypothesis that HERV mRNA expression is too low in normal tissues to achieve translation, whereas the overexpression in tumor cells is sufficient to generate epitopes efficiently presented to the immune system. Because the presence of the epitope is, at the end, the main factor required for the activity of specific T cells, it will be important to extend the immunopeptidomics analysis to tumor biopsies in the clinic [33].

Due to their viral origin, HERV-derived epitopes are expected to be highly immunogenic. As observed in our previous work in triple negative breast cancer [19], HERV-specific T cells were also detected in TILs from ovarian cancer samples. This suggests that these epitopes induced a specific T cell response in patients. Similar results have been reported for tumor epitopes derived from mutations or cancer germline antigens [39]. However, the low frequencies of specific T cells in tumors justifies the therapeutic strategy of boosting this response with a vaccine or of inducing an epitope-dependent antitumor response using T-cell based therapies. Finally, we provided evidence that HERV-specific T cells kill ovarian cancer cells in an HLA-restricted manner while sparing normal cells, in line with the immunopeptidomics data showing tumor selectivity. Because these epitopes are specific to humans, we developed an original *in vivo* model using avian embryos transplanted with ovarian tumor cells to confirm the antitumor activity of HERV-specific T cells. This model offers the possibility to use a low number of T cells and allows for the rapid establishment of the tumor with tumor-associated phenotypical features after engraftment in the embryonic tissues. The physiological relevance of this platform was demonstrated by the observation of similar response to treatment when comparing clinical data and the parallel avian tumor replicates [32,34]. In addition, it addresses the need to reduce animal use in the context of preclinical testing, as recently supported by regulatory agencies [40]. However, the lack of long-term tumor establishment and the incomplete reconstitution of the tumor stroma are limitations to take into account. Some of them are shared with the classical subcutaneous xenografts in immunocompromised mouse models used for the preclinical testing of adoptive transfer therapies (low tumor stroma complexity, poor vascularization).

In summary, we showed that HERV-K-derived epitopes are selectively presented on tumor cells in ovarian cancer. They are immunogenic in patients and induce high-avidity CD8^+^ cytotoxic T cells that selectively kill ovarian tumor cells *in vitro* and *in vivo*. These data provide the preclinical rationale for developing T-cell based therapies against these new targets, such as vaccines, adoptive T cell therapies or T cell recruiting bispecific antibodies.

## Declarations

## Ethics approval and consent to participate

NA

## Consent for publication

All authors gave their consent for publication

## Data availability statement

All data relevant to the study are included in the article or uploaded as online supplemental information.

## Acknowledgments

We wish to thank the staff of the core facilities at the Cancer Research Center of Lyon (CRCL) for technical assistance, the BRC of the CLB and Dr Olivia Le Saux for providing human samples, as well as N. Gadot and the research anatomopathology platform of the CLB. We thank T. Andrieu, P. Battiston-Montagne, and A. Jambon for assistance in flow cytometry in the CRCL Cytometry platform. The illustrations were created with https://BioRender.com. As citations were limited, we were unable to cite all available studies, and so apologize to any authors who feel their studies were not adequately represented in our accounting.

## Author contributions

Conceptualization: PB and SD. Methodology: PB, OT, YE, AP,VM, MD, SM, CD, RT, DT, EE, AV, REB, GJD, NC, NG, QW, JVG, SD

Investigation: PB, OT, YE, VM, MD, CD, RT, EE, AV, NC, QW

Supervision: JVG, SD

Writing – original draft: PB, OT, YE, SD

Writing – review & editing: PB, JVG, SD

