## Supplementary Figures and Tables for "HERV-derived epitopes represent new targets for T-cell based immunotherapies in ovarian cancer"

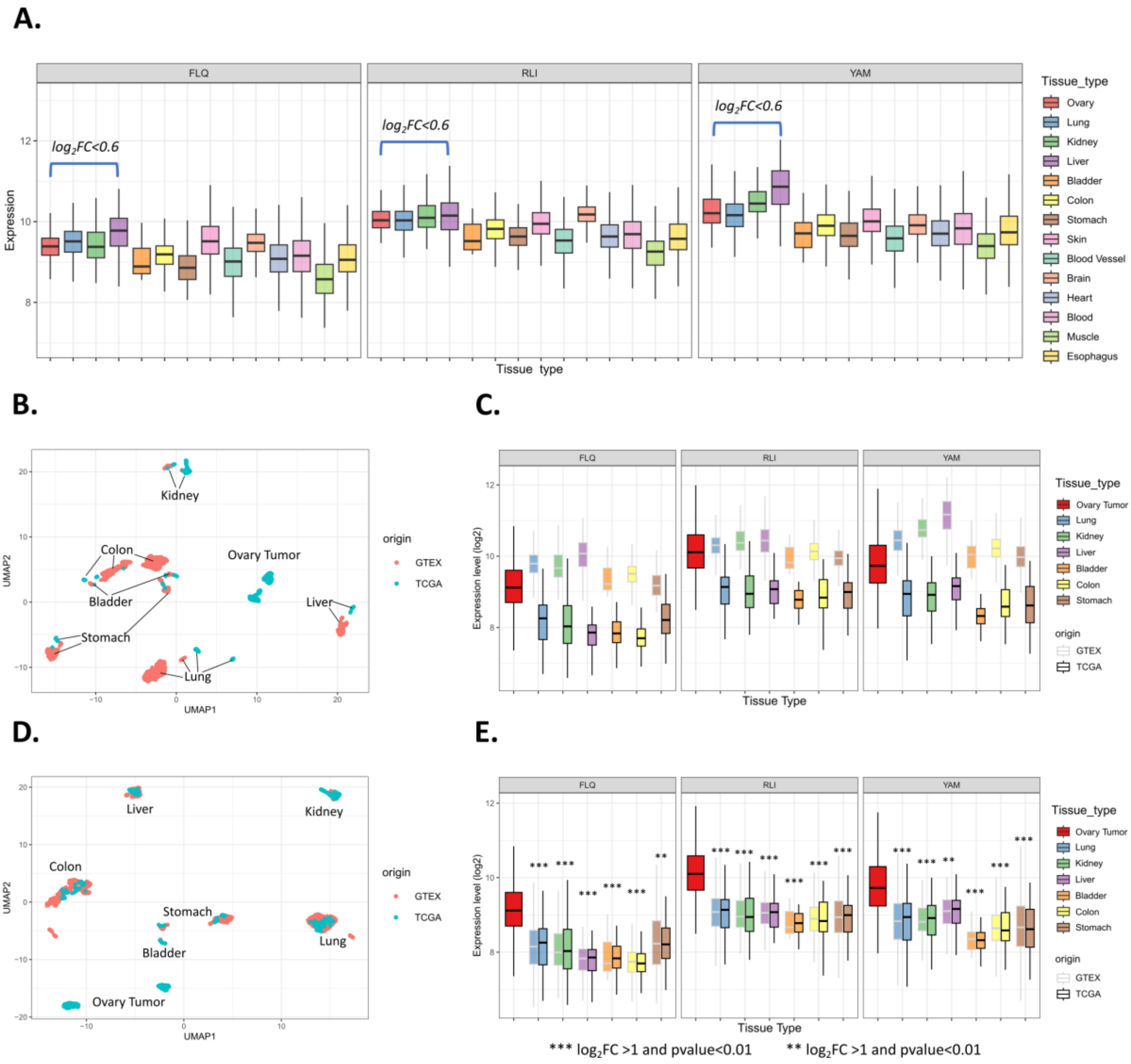


**Supplementary Figure 1: Integration of GTEx dataset to TCGA**

**A.** Epitope expression levels in normal critical tissues from GTEx. Transcriptomic expression of the epitope was estimated by the sum of all HERV loci with a potential ORF containing its sequence (FLQ, RLI, and YAM). Data were normalized and log transformed (vst) using DESeq2. The maximum positive log$_2$FC comparing healthy ovary tissue with other healthy tissues is represented for each peptide.

**B.** UMAP of merged TCGA and GTEx datasets, after normalization with DESeq2 and before any batch correction. Each dot represents a sample colored according to its dataset of origin (red: GTEx, blue: TCGA). Tissue types are highlighted nearby each cluster.

**C.** Epitope-containing HERVs expression before batch correction. Red boxes show ovary tumor samples from TCGA, the other boxes show healthy tissues from both GTEx (grey outline) and TCGA (black outline).

**D.** UMAP of merged TCGA and GTEx datasets, after normalization with DESeq2 and after batch correction with ComBat (reference batch: TCGA). Each dot represents a sample colored according to its dataset of origin (red: GTEx, blue: TCGA). Tissue types are highlighted nearby each cluster of samples.

**E.** Epitope-containing HERVs expression of merged TCGA and GTEx datasets after batch correction with ComBat. Red boxes show ovary tumor samples from TCGA, the other boxes show normal tissues from both GTEx (grey outline) and TCGA (black outline). Expression levels in ovary tumor were compared to each tissue for all 3 peptides (*** $log_2FC > 1$ and p-value<0.01, ** $log_2FC > 0.75$ and p-value<0.01, * $log_2FC > 0.5$ and p-value<0.01, Wilcoxon test).

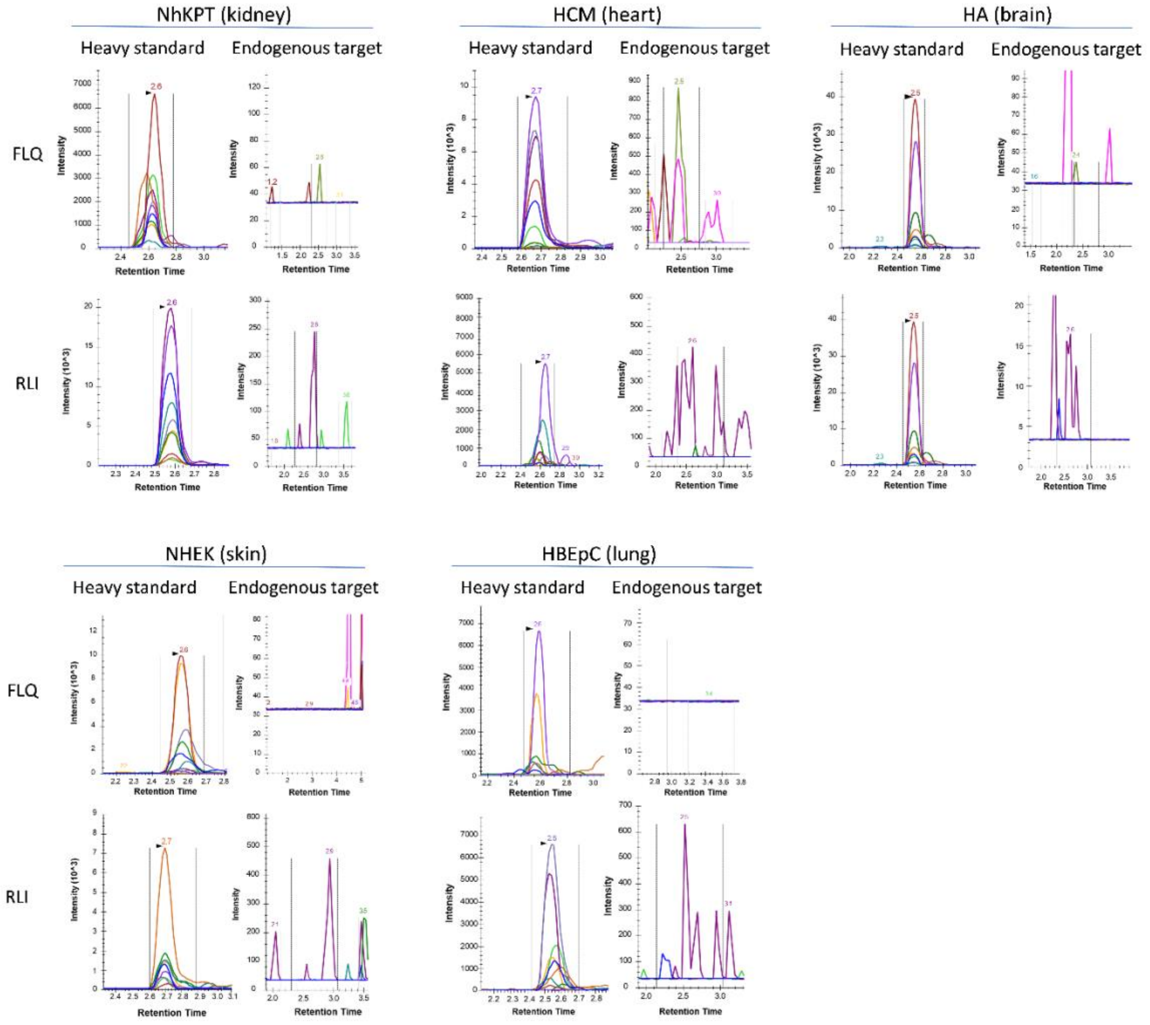


**Supplementary Figure 2: Absence of epitope presentation on normal primary cells.**

Targeted immunopeptidomics analysis (Max Rec_Ver2, Complete Omics Inc.) of FLQ and RLI presented on HLA molecules on NhKPT cells (kidney, up left), HCM (heart, up center), HA (brain, up right), NHEK (skin, down left) and HBEpC (lung, down center) primary cells**.** Heavy standard peptides are used as positive control (standard).

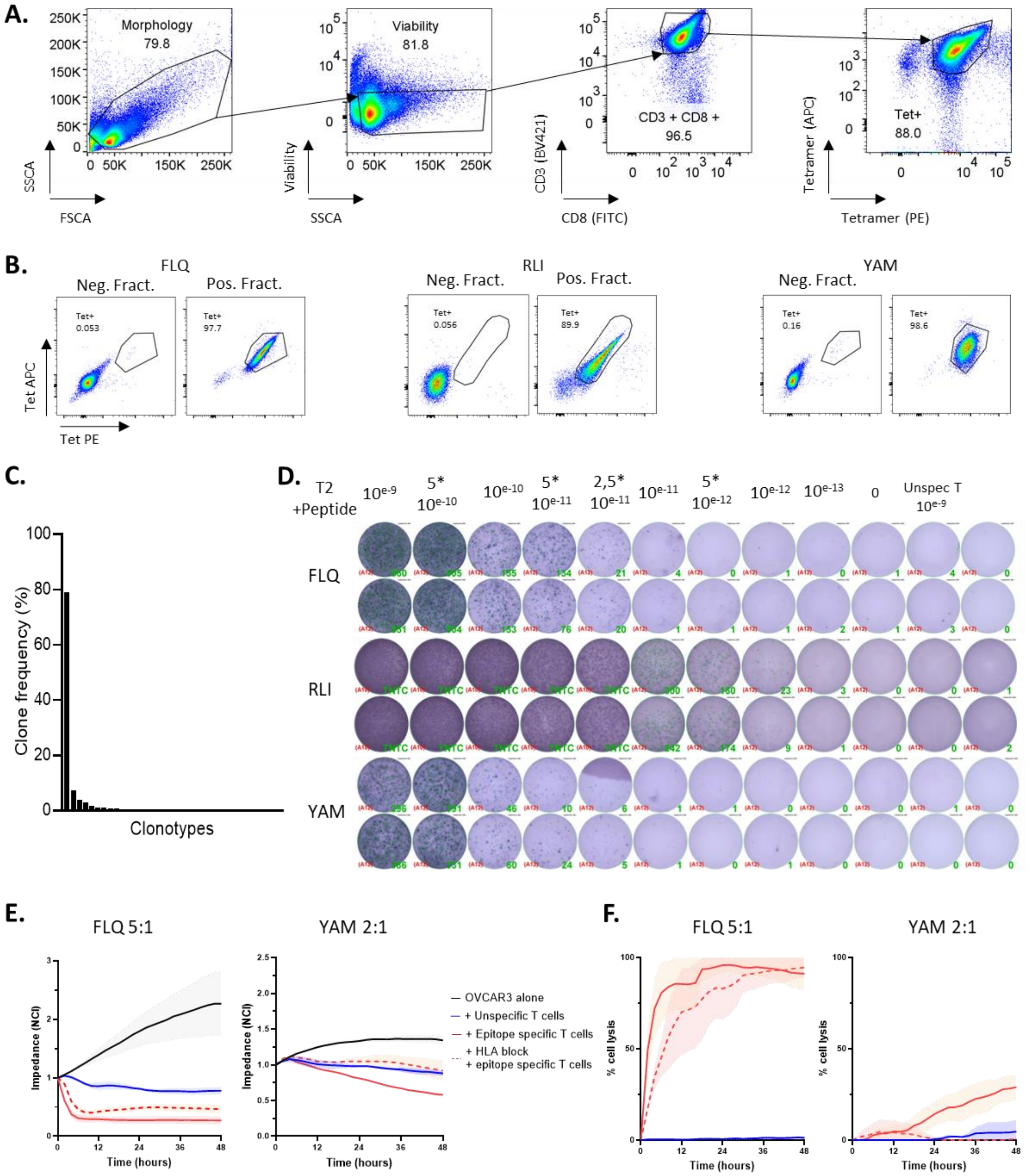


**Supplementary Figure 3: Functional analysis of HERV-K epitope-specific CD8⁺ T cells**

**A.** FACS gating strategy used to identify HERV-K epitope-specific CD8⁺ T cells.

**B.** Representative plots of FACS analysis of epitope unspecific and specific $CD8^+$ T cells. Left to right : FLQ- (n=4), RLI- (n=2) and YAM- (n=4) specific T cells. For each epitope specificity, negative and positive fractions were collected after monomer-based sorting of primed $CD8^+$ T cells.

**C.** Representative epitope (RLI)-specific $CD8^+$ T cells clonotype analysis from TCR sequencing.

**D.** Representative IFNγ ELISPOT after restimulation of tetramer-negative and positive fractions with irrelevant or cognate peptide pulsed T2 cells ($10^{-9}$ M for unspecific T cells and irrelevant peptide; $10^{-9}$ to $10^{-13}$ M range for cognate peptides).

**E and F.** Tumor cell impedance (E) and percentage of cell lysis (F) measured at each time point during co-cultures of OVCAR3 cells with FLQ- and YAM-specific T cells or tetramer-negative fractions at effector to target ratio (E:T) 5:1 (FLQ) or 2:1 (YAM). Results are mean ± s.d. of technical triplicates. Target cells alone are depicted with black lines, co-culture with specific T cells are depicted with red lines, co-culture with unspecific T cells are depicted with blue lines. Addition of the anti-HLA antibody is depicted with the red dotted line.

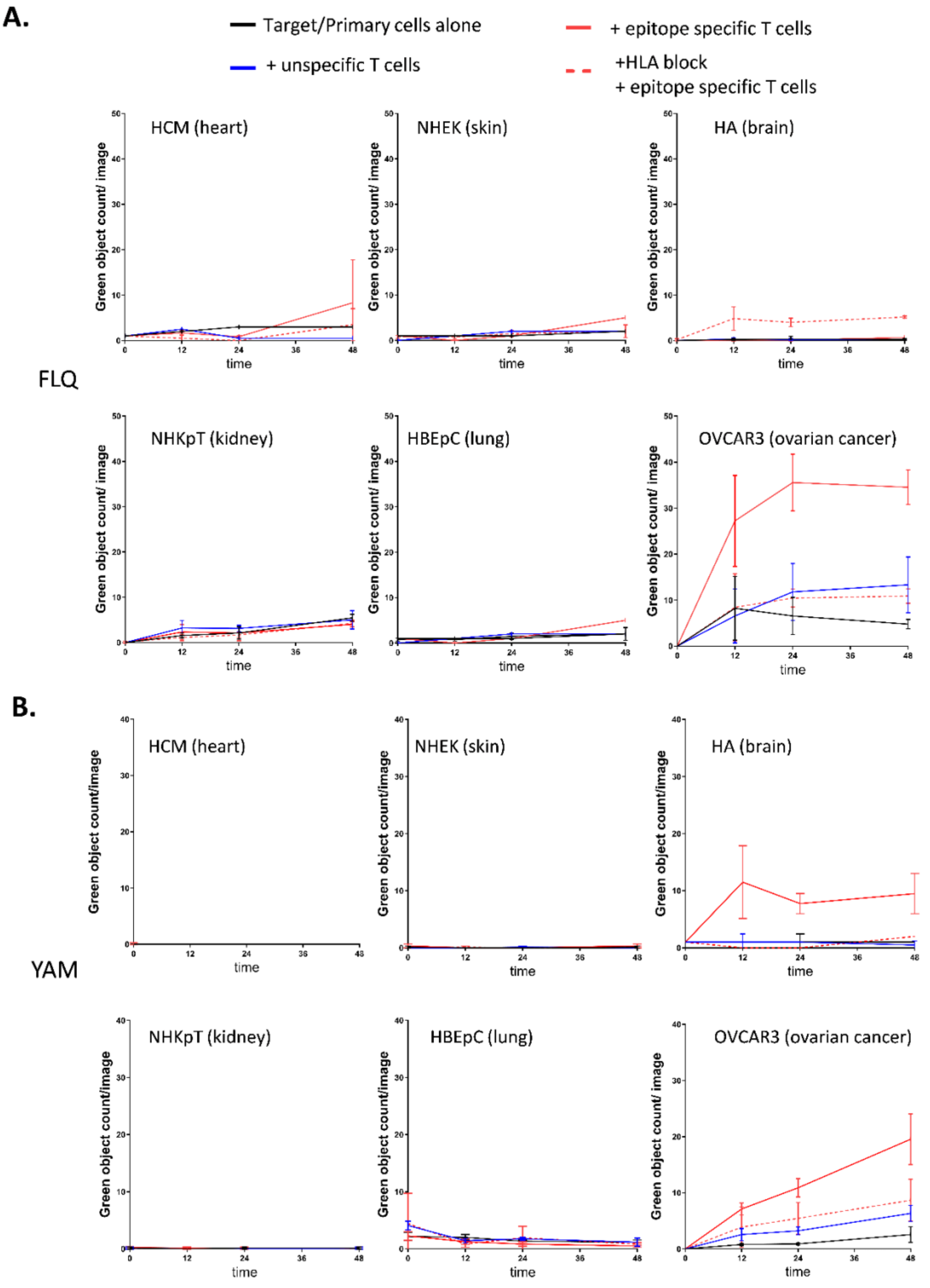


**Supplementary Figure 4: Annexin V quantification on normal primary cells after co-culture with HERV-K epitope-specific T cells**

**A.** and **B.** Annexin $V^+$ target cell death quantification at 0, 6h, 24h and 48h after the addition of FLQ- **(A)** or YAM- **(B)** specific $CD8^+$ T cells or unspecific T cells.

Target cells alone are depicted with black lines, specific T cells are depicted with red lines, unspecific T cells are depicted with blue lines. Addition of the anti-HLA antibody to specific T cells-target cells co-cultures is depicted with the red dotted line. HCM, human cardiac myocytes; NHEK, normal human epithelial keratinocytes; NHEK, normal human epithelial keratinocytes; NHKpT, normal human kidney proximal tubule cells; HBEpC, human bronchial epithelial cells.

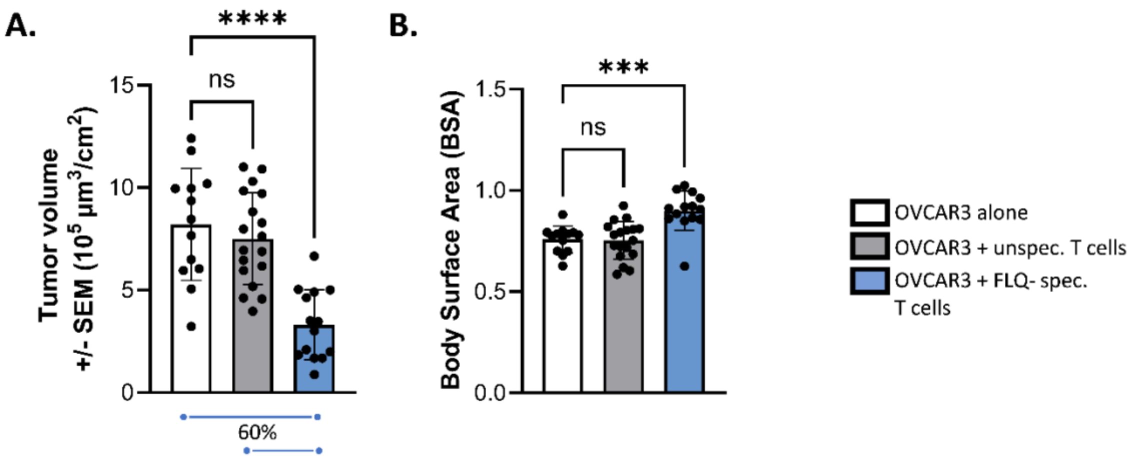


**Supplementary Figure 5: *In vivo* anti-tumoral activity of HERV-K epitope-specific CD8$^+$ T cells**

**A.** Tumor volume monitored using 3D light sheet microscopy, 48h post-engraftment.

**B**. Chicken embryo body surface areas (BSA), after 48h engraftment of OVCAR3 alone, or co-engraftment with FLQ-unspecific or specific CD8$^+$ T cells at 5:1 effector to target (E:T) ratio. Results are mean ± s.d. of embryos (OVCAR3 alone n=13; Unspecific T cells n=18; FLQ specific T cells n=14). In white OVCAR3 alone, in grey OVCAR3 and unspecific T cells co-implanted and in blue OVCAR-3 and FLQ-specific CD8$^+$ T cells co-implanted. Each point represents one embryo. **** p<0.0001, *** p<0.0003 One-way ANOVA post-hoc Dunnett's test

| FLQ | RLI | YAM |
|---|---|---|
| HML2_10q24.2 | HML2_10q24.2 | HERVK14C_13q31.1 |
| HML2_11q22.1 | HML2_11p15.4a | HERVK14C_2p12 |
| HML2_11q23.3 | HML2_11q12.3b | HML2_10p14 |
| HML2_12q14.1 | HML2_11q22.1 | HML2_10q24.2 |
| HML2_19_GL383575v2_alt | HML2_11q23.3 | HML2_11q22.1 |
| HML2_19_GL383576v1_alt | HML2_12q14.1 | HML2_19_GL383575v2_alt |
| HML2_19p12b | HML2_14q11.2 | HML2_19_GL383576v1_alt |
| HML2_19q11 | HML2_19_GL383575v2_alt | HML2_19q11 |
| HML2_1_KI270766v1_alt | HML2_19_GL383576v1_alt | HML2_19q13.12b |
| HML2_1p31.1 | HML2_19p12b | HML2_19q13.41a |
| HML2_1p36.21a | HML2_19q11 | HML2_1p36.21a |
| HML2_1p36.21b | HML2_1p31.1 | HML2_1p36.21d |
| HML2_1p36.21c | HML2_1q22 | HML2_1q22 |
| HML2_1p36.21d | HML2_1q23.3 | HML2_1q23.3 |
| HML2_1q22 | HML2_20q11.22 | HML2_20q11.22 |
| HML2_20q11.22 | HML2_22q11.21 | HML2_3p12.3 |
| HML2_22q11.21 | HML2_3p25.3 | HML2_3q13.2 |
| HML2_3q13.2 | HML2_3q21.2 | HML2_3q24 |
| HML2_3q21.2 | HML2_3q24 | HML2_3q27.2 |
| HML2_3q24 | HML2_3q27.2 | HML2_4q35.2 |
| HML2_3q27.2 | HML2_4p16.3a | HML2_5p13.3 |
| HML2_4q32.1 | HML2_4q32.1 | HML2_5q33.3 |
| HML2_5p13.3 | HML2_4q32.3 | HML2_6q14.1 |
| HML2_5q33.2 | HML2_5p12 | HML2_7p22.1 |
| HML2_5q33.3 | HML2_5p13.3 | HML2_7q22.2 |
| HML2_6q14.1 | HML2_5q33.2 | HML2_8_KI270813v1_alt(a) |
| HML2_7p22.1 | HML2_5q33.3 | HML2_8p23.1a |
| HML2_7q22.2 | HML2_6_GL000251v2_alt | HML2_8q24.3b |
| HML2_8_KI270813v1_alt(b) | HML2_6p22.1 | HML3_10q11.21a |
| HML2_8p23.1b | HML2_6q14.1 | HML3_10q22.1 |
| HML2_8q24.3a | HML2_7p22.1 | HML3_11q12.1 |
| HML2_9q34.3 | HML2_7q22.2 | HML3_12q13.12 |
| | HML2_8_KI270813v1_alt(b) | HML3_12q23.3 |
| | HML2_8p23.1b | HML3_19p12b |
| | HML2_8q24.3b | HML3_19q13.2 |
| | HML2_9q34.3 | HML3_1p21.3 |
| | | HML3_2p16.3 |
| | | HML3_4p12 |
| | | HML3_4q24a |
| | | HML3_5p13.2 |
| | | HML3_5p15.33a |
| | | HML3_6p22.1 |
| | | HML3_7p13 |

**Supplementary Table 1 - List of HERV loci containing epitope sequences.**

| No. Patient | Tissue ID. | Pathology diagnosis | Stage | HERV-K Gag Score |
|---|---|---|---|---|
| 1 | Fov020783 | High grade serous carcinoma | IA | 3 |
| 2 | Fov021200 | High grade serous carcinoma | IIA | 3 |
| 3 | Fov020833 | High grade serous carcinoma | IIA | 3 |
| 4 | Fov041285 | High grade serous carcinoma | IIB | 3 |
| 5 | Fov041143 | Mixed serous mucinous carcinoma | IIIA | 3 |
| 6 | Fov021780 | High grade serous carcinoma | IIA | 3 |
| 7 | Fov030379 | High grade serous carcinoma | IA | 2 |
| 8 | Fov020789 | High grade serous carcinoma | IIB | 2 |
| 9 | Fov040855 | High grade serous carcinoma | IIIC | 2 |
| 10 | Fov021215 | High grade serous carcinoma | IV | 2 |
| 11 | Fov021928 | Endometrioid carcinoma | IIIB | 2 |
| 12 | Fov030376 | Mucinous carcinoma | I | 2 |
| 13 | Fov040905 | Clear cell carcinoma | IIA | 2 |
| 14 | Fov040819 | High grade serous carcinoma | I | 1 |
| 15 | Fov030343 | High grade serous carcinoma | IA | 1 |
| 16 | Fov030286 | High grade serous carcinoma with necrosis | IA | 1 |
| 17 | Fov041138 | High grade serous carcinoma | III | 1 |
| 18 | Fov050836 | High grade serous carcinoma | IIIB | 1 |
| 19 | Fov030120 | High grade serous carcinoma | IIIC | 1 |
| 20 | Fov021194 | Endometrioid carcinoma | IIB | 1 |
| 21 | Fov020262 | High grade serous carcinoma | I | 0 |
| 22 | Fov030524 | High grade serous carcinoma | IA | 0 |
| 23 | Fov020386 | High grade serous carcinoma | IA | 0 |
| 24 | Fov030419 | High grade serous carcinoma | IB | 0 |
| 25 | Fov030377 | High grade serous carcinoma | IB | 0 |
| 26 | Fov030504 | High grade serous carcinoma | IC | 0 |
| 27 | Fov030124 | High grade serous carcinoma | II | 0 |
| 28 | Fov030109 | High grade serous carcinoma | II | 0 |
| 29 | Fov041120 | High grade serous carcinoma | IIB | 0 |
| 30 | Fov031171 | High grade serous carcinoma | III | 0 |
| 31 | Fov021351 | High grade serous carcinoma | IIIA1 | 0 |
| 32 | Fov050139 | High grade serous carcinoma | IIIA2 | 0 |
| 33 | Fov050700 | High grade serous carcinoma | IIIC | 0 |
| 34 | Fov040754 | High grade serous carcinoma | IIIC | 0 |
| 35 | Fov041148 | High grade serous carcinoma | IIIC | 0 |
| 36 | Fov021382 | High grade serous carcinoma | IIIC | 0 |
| 37 | Fov021786 | Endometrioid carcinoma | IIA | 0 |
| 38 | Fov021360 | Endometrioid carcinoma | IIB | 0 |
| 39 | Fov040968 | Mucinous carcinoma | IIA | 0 |
| 40 | Fov040870 | Mucinous carcinoma | IIA | 0 |

**Supplementary Table 2 – Tumor microarrays, patients' information.**

Patient informations include tissue ID, pathology diagnosis, stage and IHC scoring.

Patient were classified according to the HERV-K/HML-2 Gag IHC score;  n=31 high-grade serous carcinoma, n=1 mixed carcinoma, n=1 clear cell carcinoma, n=4 endometrioid carcinoma.
